# Eukaryotic Acquisition of a Bacterial Operon

**DOI:** 10.1101/399394

**Authors:** Jacek Kominek, Drew T. Doering, Dana A. Opulente, Xing-Xing Shen, Xiaofan Zhou, Jeremy De Virgilio, Amanda B. Hulfachor, Cletus P. Kurtzman, Antonis Rokas, Chris Todd Hittinger

## Abstract

Operons are a hallmark of bacterial genomes, where they allow concerted expression of multiple functionally related genes as single polycistronic transcripts. They are rare in eukaryotes, where each gene usually drives expression of its own independent messenger RNAs. Here we report the horizontal operon transfer of a catecholate-class siderophore biosynthesis pathway from Enterobacteriaceae into a group of closely related yeast taxa. We further show that the co-linearly arranged secondary metabolism genes are actively expressed, exhibit mainly eukaryotic transcriptional features, and enable the sequestration and uptake of iron. After transfer to the eukaryotic host, several genetic changes occurred, including the acquisition of polyadenylation sites, structural rearrangements, integration of eukaryotic genes, and secondary loss in some lineages. We conclude that the operon genes were likely captured in the shared insect gut habitat, modified for eukaryotic gene expression, and maintained by selection to adapt to the highly-competitive, iron-limited environment.

## Main Text

The core processes of the Central Dogma of Biology, transcription and translation, are broadly conserved across living organisms. Nonetheless, there are seemingly fundamental differences between the domains of life in how these processes are realized. Eukaryotic transcription is spatially and temporally separated from translation and generally operates on individual genes through a complex interplay of transcription factors and chromatin remodeling complexes. Nascent mRNAs are co-transcriptionally processed by adding 3’ polyadenosine (poly(A)) tails and 5’ caps of 7-methyl-guanosine (m^7^G) before they are trafficked out of the nucleus for translation. In bacteria, transcription is tightly coupled with translation, and both occur inside the cytosol. Furthermore, bacterial transcription often operates on clusters of genes, known as operons, where a single regulatory region regulates the expression of physically-linked genes into a polycistronic mRNA that is minimally processed and translated into several polypeptides at similar abundance. In contrast, eukaryotic operons, which are rare in most taxa but are frequently found in nematodes (*1*, *2*) and tunicates (*3*, *4*), are processed by trans-splicing and related mechanisms. Operon dissemination has been proposed to occur predominantly via horizontal gene transfer (HGT) (*5*, *6*), a process where organisms acquire genes from sources other than their parents. HGT is pervasive and richly documented among bacteria, but it is thought to be rarer in eukaryotes (*7*–*11*). Known examples of bacterium-to-eukaryote HGT occurred as single genes, but never as operons. Nonetheless, horizontal operon transfer (HOT) into eukaryotes would allow even complex pathways to spread rapidly, especially in environments where competition for key nutrients is intense.

One such nutrient is iron, which plays crucial roles in many essential cellular processes (*12*–*14*) and is a key determinant of virulence in both animal and plant pathogens (*15*–*17*). Many specialized systems have evolved to sequester it from the surrounding environment, one of which is the biosynthesis of small-molecule iron chelators called siderophores. Most bacteria synthesize catecholate-class siderophores (*18*), whereas hydroxamate-class siderophores are commonplace in fungi (*19*). A notable exception is the budding yeast lineage (subphylum Saccharomycotina), which has long been thought to completely lack the ability to synthesize their own siderophores, despite its ability to utilize them (*19*). Here we survey a broad range of fungal genomes for known components of iron uptake and storage systems. Although most systems are broadly conserved, we identify a clade of closely related yeast species that contains a bacterial siderophore biosynthesis pathway. Through phylogenetic hypothesis testing, we show that this pathway was acquired through horizontal operon transfer (HOT) from the bacterial family Enterobacteriaceae, which includes *Escherichia coli, Erwinia carotovora, Yersinia pestis*, and relatives that share the insect gut niche with many of these yeasts (*20*). After acquisition, the operon underwent structural changes and progressively gained eukaryotic characteristics, while maintaining the clustering of functionally related genes. Transcriptomic experiments and analyses show that the siderophore biosynthesis genes are actively expressed, contain poly(A) tails, and exhibit evidence of mostly monocistronic transcripts, as well as some potentially bicistronic transcripts. *In vivo* assays also demonstrate the biosynthesis and secretion of functional catecholate-class siderophores in several of these yeast species. This remarkable example shows how eukaryotes can acquire a functional bacterial operon, while modifying its transcription to domesticate and maintain expression as a set of linked eukaryotic genes.

## Results

### Iron uptake and storage is conserved in fungi

We surveyed the genome sequences of 175 fungal species and observed broad conservation of genes involved in low-affinity iron uptake, vacuolar iron storage, reductive iron assimilation, heme degradation, and siderophore import systems (Fig. 1, Table S1). In contrast, genes involved in siderophore biosynthesis pathways were more dynamic. Siderophore biosynthesis was thought to be completely absent in budding yeasts (*19*), but the genomes of *Lipomyces starkeyi* and *Tortispora caseinolytica* contain homologs of the *SidA, SidC, SidD, SidF,* and *SidL* genes involved in the biosynthesis of ferricrocin and fusarinine C, which are hydroxamate-class siderophores synthesized from L-ornithine by many filamentous fungi, such as *Aspergillus nidulans* (*19*). Since these species are the earliest-branching budding yeast taxa, the presence of this pathway in their genomes is likely an ancestral trait inherited from the last common ancestor of the Pezizomycotina and Saccharomycotina, while its absence in most yeasts is likely due to a loss early in budding yeast evolution. Surprisingly, the genomes of three closely related Trichomonascaceae species (*Candida versatilis, Candida apicola,* and *Starmerella bombicola*) contain multiple homologs of bacterial siderophore biosynthesis genes (*entA-F*) that are predicated to enable the synthesis of catecholate-class siderophores from chorismate (*21*) (Fig. S1). These genes are co-linear and predicted to be expressed from the same strand of DNA, features that are both reminiscent of the operons where these genes are found in bacteria.

**Fig. 1.**
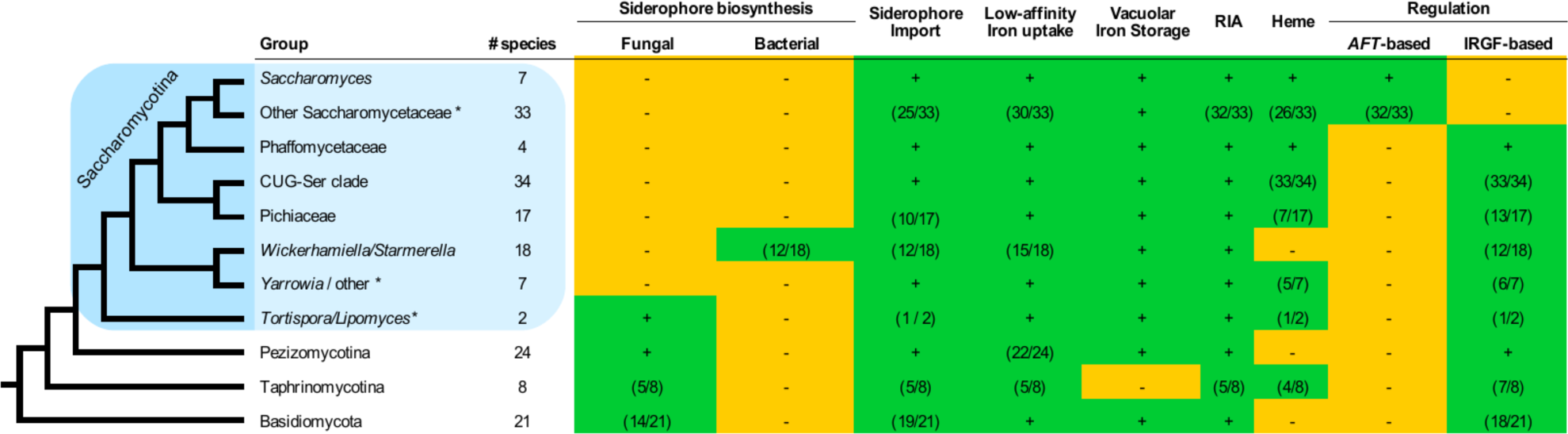
Distribution of the iron uptake and storage systems among fungi. Plus (green) and minus (orange) signs indicate the presence and absence of iron uptake and storage systems in specific taxonomic groups. The numbers in parentheses (green) indicate the number of species in a taxonomic group that possess a specific system, if it is not ubiquitous in that group. Blue box indicates the budding yeasts. RIA - Reductive Iron Assimilation. IRGF – Iron-Responsive GATA Factor. For details about specific taxa and individual genes see Table S2. Asterisks (*) mark paraphyletic groups. Note that only *Wickerhamiella/Starmerella* (W/S) clade fungi contain the bacterial or catecholate-class siderophore biosynthesis pathway.

### Horizontal operon transfer (HOT) from bacteria to yeasts

To investigate the evolutionary history of these genes, we sequenced and analyzed 18 additional genomes from the *Wickerhamiella/Starmerella* clade (W/S clade, Table S2) and identified the catecholate-class siderophore biosynthesis pathway in 12 of these species (Fig. 2a, 2c). To determine whether the yeast siderophore biosynthesis genes were horizontally acquired from a bacterial operon, we first used the *ent* genes found in yeasts to perform BLAST queries against the bacterial data present in GenBank and found that the top hits belonged to a range of species from the family Enterobacteriaceae. Since no single taxon was overrepresented, we surveyed 1,336 publicly available genomes from the class Gammaproteobacteria, to which the Enterobacteriaceae belongs, for the presence of *entA-entF* homologs and extracted them from all 207 genomes where all six genes could be reliably identified (Table S3). We then reconstructed unconstrained maximum-likelihood (ML) phylogenies for each *ent* gene, as well as for a concatenated super-alignment of all six genes (*entABCDEF,* Table S4). Since *entF* contributed nearly two-thirds of the total alignment length, we also evaluated a super-alignment of the remaining five genes (*entABCDE,* Fig. 2a).

**Fig. 2.**
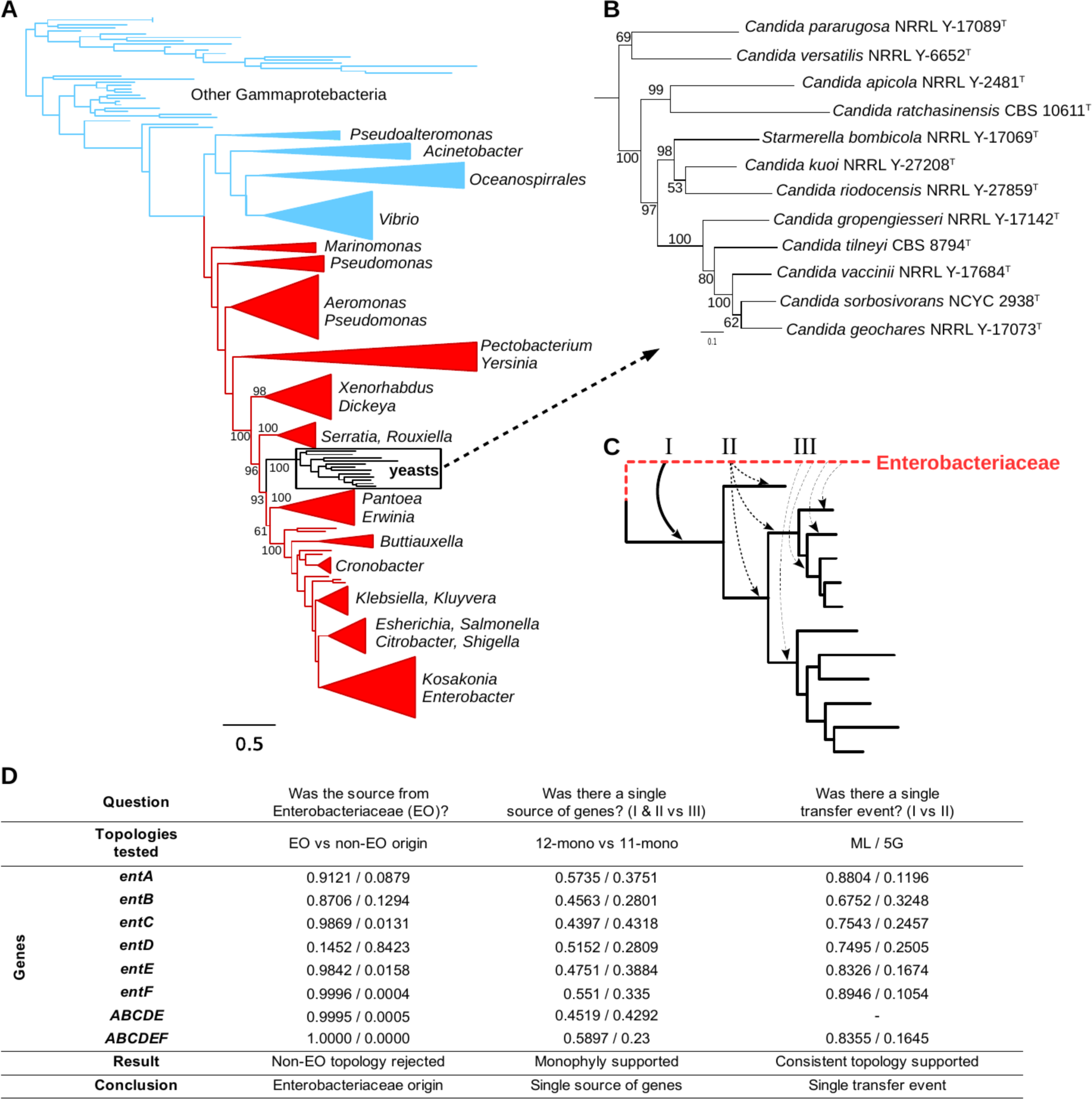
Yeast siderophore biosynthesis originated from an Enterobacteriaceae lineage. (B) ML phylogeny from the super-alignment of *entABCDE* genes from 207 Gammaproteobacteria and 12 yeasts, rooted at the midpoint. Bootstrap support values are shown for relevant branches within the Enterobacteriaceae (red). Other Gammaproteobacteria are green. (B) Detailed view of the yeast clade from the main phylogeny, with bootstrap supports. (C) Alternative scenarios for the horizontal operon transfer. (D) P-values of the AU test of different evolutionary hypotheses tested in this study; EO – Enterobacteriaceae origin; non-EO – non-Enterobacteriaceae origin; 12-mono - 12 yeast sequences are monophyletic, 11-mono - 11 yeast sequences monophyletic and one unconstrained (12 alternatives tested, lowest p-value shown, full details in Table S5); 5G – topology of the yeast clade constrained to the one inferred from the super-alignment of *entABCDE* genes.

**Fig. 3.**
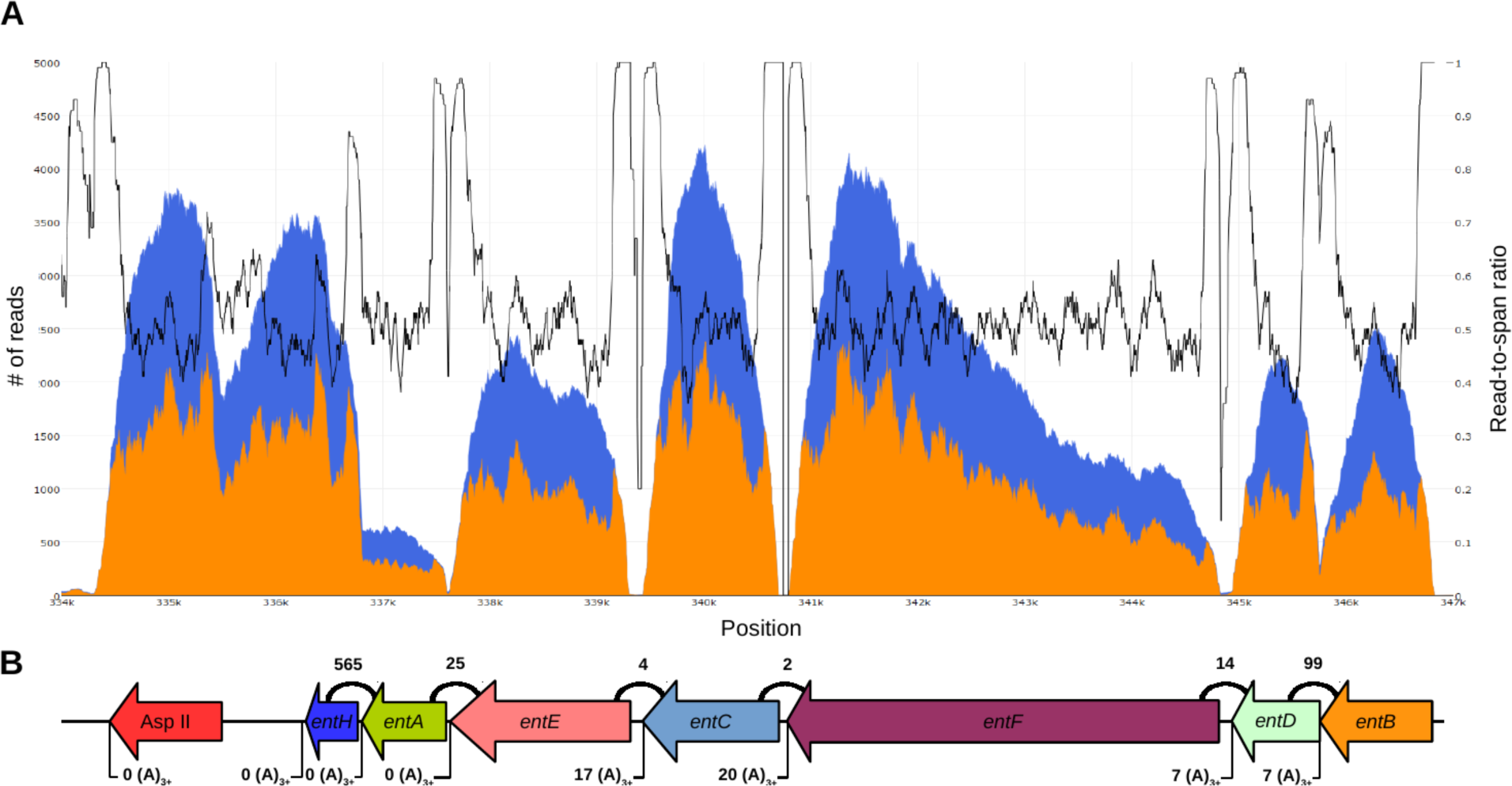
Transcriptomics of the siderophore biosynthesis genes in *C. versatilis.* The orange area indicates per-base coverage by RNA-seq reads (read coverage). The blue area indicates per-base cumulative coverage by RNA-seq reads and inserts between read pairs (span coverage). The black line indicates the ratio of the read coverage over the span coverage, which is expected to remain ∼50% in the middle of gene transcripts and rise towards 100% at transcript termini. The expected 3’ coverage bias can be observed for individual transcripts in the raw coverage data. Diagram of siderophore biosynthesis genes in the *C. versatilis* genome, drawn to scale, as well as a gene encoding a class II asparaginase adjacent on the 3’ end. Counts above indicate read pairs cross-mapping between genes. Counts below indicate reads containing putative poly(A) tails.

Consistent with the BLAST results, the yeast sequences formed a highly-supported, monophyletic group nested within the Enterobacteriaceae lineage on all gene trees, placing their donor lineage after the divergence of the *Serratia/Rouxiella* lineage and before the divergence of the *Pantoea/Erwinia* lineage from closer relatives of *E. coli*. To formally test the hypothesis of an Enterobacteriaceae origin, we reconstructed phylogenies under the constraints that yeast sequences either group together with the Enterobacteriaceae (EO) or outside of that clade (non-EO). We then employed the approximately unbiased (AU) test to determine if the EO phylogenies were a statistically better explanation of the data than the non-EO phylogenies. The EO phylogeny was strongly preferred (p-value < 10^−3^) for the six- and five-gene concatenation data matrices (Fig. 2d). Individual genes carried weak signal due to their short lengths, but the *entC, entE,* and *entF* genes nonetheless strongly supported the Enterobacteriacae origin (p-value < 0.05), *entA* and *entB* had consistent but weaker support, and no individual gene rejected the EO hypothesis. Next, we sought to determine the course of the transfer event and tested a single-source, single-transfer hypothesis against multi-source and multi-transfer alternatives, each of which predicted specific phylogenetic patterns (Fig. 2c). AU tests on the reconstructed phylogenies did not support multiple transfer events and instead supported the simplest explanation that the HOT event occurred from a single source lineage directly into a single common ancestor of the W/S clade yeasts (Fig. 2d).

### Transferred genes have mainly eukaryotic transcript features

To determine whether and how these yeasts overcame the differences between eukaryotic and bacterial gene expression, we sequenced mRNA from *C. versatilis, C. apicola,* and *St. bombicola*. These species were chosen due to their diverse gene cluster structures and positions on the phylogenetic tree: *C. versatilis* was chosen as an early-branching representative whose structure appeared to be more similar to the ancestral operon, while *St. bombicola* and *C. apicola* appeared to represent more derived stages of evolution in the eukaryotic hosts. Each of the three species expressed mRNAs for the siderophore biosynthesis genes, and *C. versatilis* expression was the highest (Table S6).

We then examined the transcriptomic data for characteristics that are typically bacterial or eukaryotic. The length of intergenic regions was not divisible by three, so we immediately excluded the hypothesis that they were translated as a single fused polypeptide. The *C. versatilis* genes were expressed at similar levels, whereas *St. bombicola* and *C. apicola* genes showed significant diversity in their expression (Table S6, Fig. 4, Figs S2-S4). Interestingly, we also observed that the siderophore biosynthesis genes in *C. versatilis* had much shorter intergenic sequences than their counterparts in *St. bombicola* and *C. apicola,* which were each shorter than their respective genome-wide means (within gene cluster intergenic means were 158, 484, and 377 bps versus genome-wide means of 370, 549, and 455 bps for *C. versatilis, St. bombicola* and *C. apicola*, respectively). Shorter intergenic distances can enhance transcriptional coupling between neighboring genes inside operons (*22*, *23*), so these results suggest that *C. versatilis* might have retained this feature due to selection for concerted expression.

**Fig. 4.**
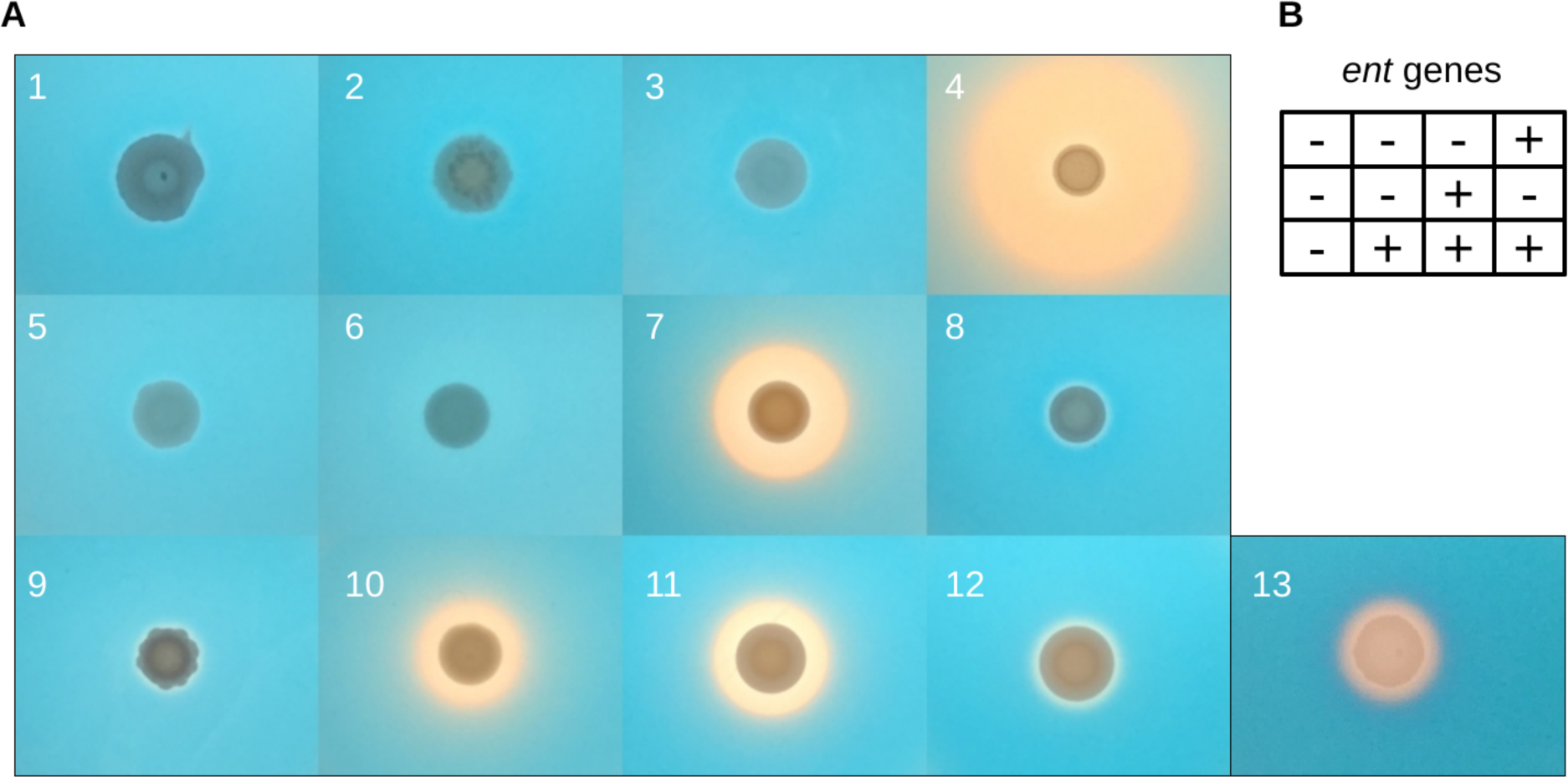
Siderophore production by yeasts from the *Starmerella/Wickerhamiella* clade. (A) CAS-based overlay assay of siderophore production. Under normal conditions, the medium remains blue, but in the presence of an iron chelator, it changes color from blue to orange. Species legend: (1) *Saccharomyces cerevisiae* FM1282, (2) *Yarrowia lipolytica* NRRL YB-423^T^, (3) *Candida hasegawae*, (4) *Candida pararugosa,* (5) *Wickerhamiella cacticola*, (6) *Wickerhamiella domercqiae*, (7) *Candida versatilis*, (8) *Candida davenportii*, (9) *Candida apicola*, (10) *Candida riodocensis*, (11) *Candida kuoi*, (12) *Starmerella bombicola*, (13) *Escherichia coli* MG1655 (positive control). Results of the CAS assay for all analyzed species can be found in Fig. S5. (B) Distribution of siderophore biosynthesis genes in the genomes of species depicted in panel A.

To further investigate operon-like characteristics that may have been retained, we analyzed read pairs in which the forward and reverse reads mapped to different genes, providing physical evidence of transcripts composed of multiple genes. To quantify this signal, we calculated the per-site ratio of the actual sequence coverage and the coverage spanned by the inserts between read pairs (i.e. coverage/span coverage). Ratios of 50% are expected for most of the length of a transcript, while ratios of 100% indicate the ends of the transcripts. Thus, transcript boundaries are visualized as a coverage trough between two spikes approaching 100% ratios. Ratios below 100% at the putative 5’ or 3’ ends of annotated transcripts, coupled with non-zero coverage of their intergenic regions, provide evidence of overlapping (and potentially bicistronic) transcripts. Most transcripts predicted to be involved in siderophore biosynthesis were monocistronic in *St. bombicola, C. apicola,* and *C. versatilis,* but *C. versatilis* had a sub-population of potentially overlapping mRNAs, including the *entB* and *entD* genes on one end, as well as the *entE, entA,* and *entH* genes on the other (Fig. 4), with the *entE-entA-entH* genes showing the strongest signal of overlap. Previously reported yeast bicistronic transcripts have been attributed mainly to inefficiencies in the RNA transcription machinery (*24*, *25*), whereas the yeast *ent* transcripts we have described here encode functionally related steps of a biosynthesis pathway that may retain some polycistronic characteristics from their ancestry as parts of a bacterial operon.

We also examined the transcriptomic data for evidence of transcriptional processing and found that many of the siderophore biosynthesis genes contained putative polyA tails (Fig 4., Figs S2-S4). We did not find any evidence suggesting that 5’ caps were added by trans-splicing (*26*) or by alternatively cis-splicing a common cassette exon upstream of each protein-coding region (*27*). Thus, we conclude that, even in *C. versatilis*, the majority of transcripts are likely transcribed and processed through conventional eukaryotic mechanisms that involve distinct promoters and polyadenylation sites for each gene. These results further suggest that most sequence modifications for eukaryotic expression act pre- or co-transcriptionally, rather than through specialized sequences to enable translation.

### Bacterial siderophore biosynthesis is functional in yeasts

To determine whether yeasts that contain the *ent* biosynthesis genes actually produce siderophores, we grew them on a low-iron medium overlaid with iron-complexed indicators. In presence of iron chelators, such as siderophores, the medium changes color from blue to orange, in a characteristic halo pattern that tracks the diffusion gradient of siderophores secreted from colonies into the surrounding medium. We tested the 18 yeast species from the W/S clade that we sequenced, together with eight outgroup species spread broadly across the yeast phylogeny (including *S. cerevisiae*) and *E. coli* as a positive control, and we observed unambiguously strong signals of siderophore production in five species, all of which contained the siderophore biosynthesis genes (Fig. 4, Fig. S5). The lack of signal in other species harboring the siderophore biosynthesis genes could suggest the secondary inactivation of the pathway (through mechanisms other than nonsense or frameshift mutations, which are absent), but it is more likely that siderophore production is below the sensitivity of the CAS assay or is not induced under the conditions studied. Nevertheless, this experiment conclusively shows that the bacterial siderophore biosynthesis are, not only transcriptionally active, but also fully functional in at least some W/S clade yeasts.

### Evolution of a bacterial operon inside a eukaryotic host

Given the significant differences in Central Dogma processes between bacteria and eukaryotes, we investigated how the horizontally transferred operon was successfully assimilated into these yeasts by mapping key changes in gene content, structure, and regulation onto the phylogeny (Fig. 5a). First, the phylogenetic distribution of the operon genes suggests at least five cases of secondary loss in W/S clade yeasts, a common occurrence for other fungal gene clusters (*28*–*31*). Although all taxa contain the six core genes (*entA-F*), *C. versatilis* uniquely harbors a homolog of the *entH* gene, which encodes a proofreading thioesterase that is not strictly required for siderophore biosynthesis (*32*). Since no homologs or remnants of other genes from the bacterial operon could be identified, we hypothesize that they were lost due to functional redundancy with genes already present in yeast genomes (e.g. the bacterial ABC transporters *fepA-G* are redundant with the yeast major facilitator superfamily transporters *ARN1-4*, the bacterial esterase *fes* is redundant with yeast ferric reductases *FRE1-8*). Second, most extant Enterobacteriaceae species closely related to the source lineage share an operon structure similar to that of *E. coli* (Table S4), which is more complex than that of the W/S clade yeasts (Fig. 5b). Based on this evidence and a molecular clock (*33*), we infer that an ancient bacterial operon, whose structure was somewhere between that of *E. coli* and *C. versatilis*, was horizontally transferred into a yeast cell tens of millions of years ago. The operon may have contained fewer genes than extant bacterial operons, or some shared gene losses or rearrangements may have occurred to produce a structure similar to that of *C. versatilis* in the last common ancestor of the W/S clade yeasts. Modern yeasts of this clade evolved at least four different structures through several lineage-specific rearrangements that tended to create derived gene cluster structures with more eukaryotic characteristics, including increasing the size of the intergenic regions, splitting the gene cluster in two in *C. apicola*, and intercalating at least four eukaryotic genes. The intercalation of a gene encoding a eukaryotic ferric reductase (*FRE*), which is involved in reductive iron assimilation, between two operon genes in a subset of species offers a particularly telling example. The genetic linkage of these two mechanisms for acquiring iron shows that bacterial and eukaryotic genes can stably co-exist, and perhaps even be selected together as gene clusters for co-inheritance or co-regulation, through eukaryotic mechanisms.

**Fig. 5.**
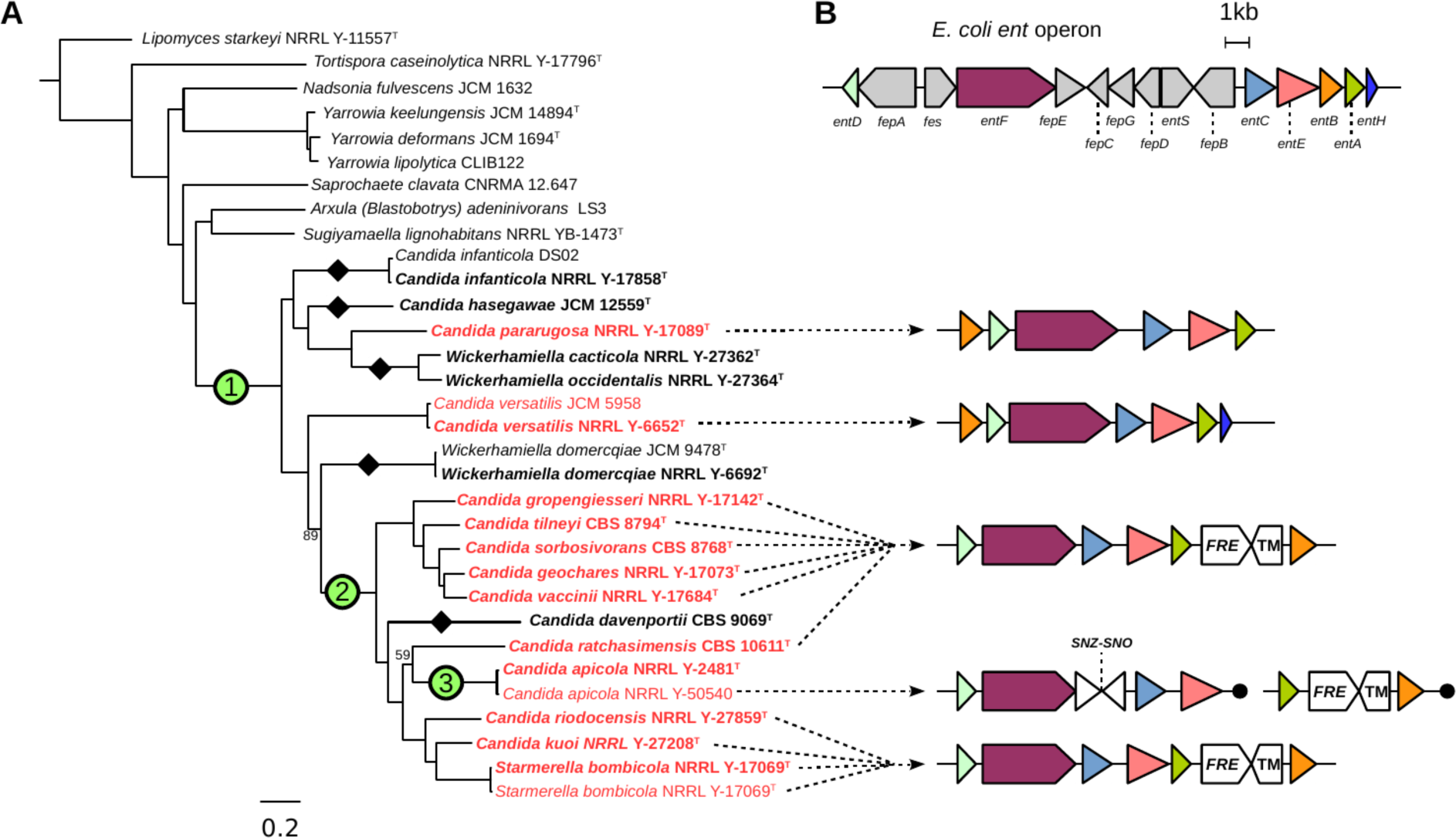
Evolution of the siderophore biosynthesis genes in yeasts. (A) ML phylogeny reconstructed from the concatenated alignment of 661 conserved, single copy genes (834,750 sites), with branch supports below 100 shown. Species in bold denote genomes sequenced in this study, while species in red denote genomes containing the siderophore biosynthesis genes. Black diamonds indicate secondary losses in yeast lineages, accompanied by losses of the siderophore importer *ARN* genes, which are often found in close proximity. (1) Horizontal operon transfer from an Enterobacteriaceae lineage. (2) Rearrangement and integration of genes encoding ferric reductase (*FRE*) and an uncharacterized transmembrane protein (TM). (3) Disruption by integration of the *SNZ-SNO* gene pair and translocation. (B) Genetic structure of the siderophore biosynthesis operon in *E. coli* and yeasts. Individual colors represent homologous ORFs, drawn to scale, and gray marks genes not found in yeasts. Black circles represent contig termini within 25kb.

### Discussion

The horizontal transfer of this siderophore biosynthesis operon is the first clearly documented example of the acquisition of a bacterial operon by a eukaryotic lineage. Several examples of horizontal gene transfer between different domains of life have been uncovered (*9*, *34*–*37*), but the transfer of entire operons into eukaryotes has been merely speculated upon as an intriguing potential route of acquisition of secondary metabolism pathways (*34*, *38*). The previous lack of evidence for HOT into eukaryotes led authors to propose barriers due to pathway complexity (*39*) and differences in core Central Dogma processes (*7*, *40*). Where could the transfer of the siderophore biosynthesis operon between Enterobacteriaceae and yeasts have occurred, and how could the bacterial operon have been functionally maintained in the yeasts’ genomes? Eukaryotes have been proposed to acquire bacterial genes through several mechanisms, including virus-aided transmission (*41*), environmental stress-induced DNA damage and repair (*42*, *43*), and a phagocytosis-based gene ratchet (*44*). The species that harbor the siderophore biosynthesis operon have been isolated predominantly from insects (*45*–*47*), where stable bacterial and eukaryotic communities coexist inside their guts (*20*). Moreover, this niche harbors diverse Enterobacteriaceae populations in which horizontal gene transfer has been reported (*48*, *49*), and insect guts have recently been described as a “mating nest” for yeasts (*50*). Since Enterobacteriaceae and yeasts can conjugate directly in some cases (*51*), it is plausible that the last common ancestor of the W/S clade yeasts incorporated the operon from a bacterial co-inhabitant of an insect gut. Due to the intense competition for nutrients in this ecosystem, including a constant arms race with the host organism itself (*52*), yeasts able to make their own siderophores and sequester iron may have had a substantial advantage over those relying on siderophores produced by others.

Given the fundamental differences between bacterial and eukaryotic gene regulation, how could a bacterial operon have been maintained in a eukaryotic genome upon transfer? If it had not been actively expressed and functional, the genes of the operon would have been rapidly lost from the genome through neutral evolutionary processes. Although eukaryotes do not encode proteins with significant similarity to the bacterial regulator Fur that controls the expression of the bacterial *ent* genes, their iron response is governed by transcription factors that also belong to the GATA family. Indeed, the consensus Fur-binding site (5’-GATAAT-3’) is remarkably similar to that of the fungal transcriptional factors that respond to iron (5’-WGATAA-3’) (*19*, *53*). This similarity suggests the intriguing possibility that the siderophore genes could have readily switched from being regulated by a bacterial transcription factor to a eukaryotic transcription factor, at least for the most 5’ promoter. Siderophores are potent chelators that can efficiently sequester iron even at low concentrations (*54*), so even a low basal expression level of the newly acquired bacterial genes could have been enough to convey a considerable selective advantage. This initial eukaryotic expression, perhaps aided by noisy transcriptional and translation processes that include leaky scanning and internal ribosome entry sites (IRESs), could then have been optimized by acquiring more eukaryotic characteristics, such as longer intergenic regions that were gradually refined into promoters, distinct polyadenylation sites, and a shift from polycistronic to bicistronic and eventually to primarily monocistronic transcripts. The incorporation of a eukaryotic gene encoding a ferric reductase would have further improved the efficiency of iron acquisition in the highly competitive ecological niche of insect guts, while enhancing the eukaryotic characteristics of the gene cluster. Our HOT finding dramatically expands the boundaries of the cross-domain gene flow. The transfer, maintenance, expression, and adaptation of a multi-gene bacterial operon to a eukaryotic host underscore the flexibility of transcriptional and translational systems to produce adaptive changes from novel and unexpected sources of genetic information.

## Acknowledgments

We thank David J. Eide and Michael D. Bucci for advice on low-iron media; Nicole T. Perna and Jeremy D. Glasner for *E. coli* strain MG1655; the Eide, Perna, Rokas and Hittinger labs for comments and discussions; RIKEN for publicly releasing 20 genome sequences; Lucigen Corporation (Middleton, WI) for use of their Covaris for gDNA sonication; and the University of Wisconsin Biotechnology Center DNA Sequencing Facility for providing Illumina sequencing facilities and services. This work was conducted in part using the computational resources of the Wisconsin Energy Institute and the Center for High-Throughput Computing at the University of Wisconsin-Madison. This material is based upon work supported by the National Science Foundation under Grant Nos. DEB-1442113 (to A.R.) and DEB-1442148 (to C.T.H. and C.P.K.), in part by the DOE Great Lakes Bioenergy Research Center (DOE Office of Science BER DE-FC02-07ER64494 to Timothy J. Donohue), the USDA National Institute of Food and Agriculture (Hatch Project 1003258 to C.T.H.), and the National Institutes of Health (NIAID AI105619 to A.R.). C.T.H. is a Pew Scholar in the Biomedical Sciences, supported by the Pew Charitable Trusts. D.T.D. is supported by a NHGRI training grant to the Genomic Sciences Training Program (5T32HG002760). Mention of trade names or commercial products in this publication is solely for the purpose of providing specific information and does not imply recommendation or endorsement by the U.S. Department of Agriculture. USDA is an equal opportunity provider and employer.

Raw DNA and RNA sequencing data were deposited in GenBank under Bioproject ID PRJNA396763. Whole Genome Shotgun assemblies have been deposited at DDBJ/ENA/GenBank under the accessions NRDR00000000-NREI00000000. The versions described in this paper are versions NRDR01000000-NREI01000000.

Author contributions: J.K. (study design, genome assembly, annotation, phylogenetic analyses, RNA-seq data analysis, text); D.T.D. (study design, CAS assays, RNA isolation and strand-specific library preparation, text); D.A.O., J.D., and A.B.H. (genomic DNA isolation and library preparation); X.S and X.Z. (preliminary genomic analyses); and C. P. K., A.R., and C.T.H. (study design, text).

## List of Supplementary Materials

Materials and Methods

Figures S1-S5

Captions for Tables S1-S6 (separate files)

## Supplementary Materials

### Materials and Methods

#### Identification of genes involved in iron uptake and storage

Amino acid sequences of proteins known to be involved in iron uptake and storage were used as BLASTP and TBLASTN v2.2.28+ (*55*) queries against genomes and proteomes of a broad range of fungal species (see Table S1 for complete list of proteins and genomes). The genomic data was obtained from GenBbank as well as from draft genome assemblies generated for 20 strains by the RIKEN BioResource Center and RIKEN Center for Life Science Technologies through the Genome Information Upgrading Program of the National Bio-Resource Project of the MEXT. *S. cerevisiae* homologs were used, except for the fungal hydroxamate-class siderophore biosynthesis proteins, which came from *A. nidulans;* the bacterial catecholate-class siderophore biosynthesis proteins, which came from *E. coli;* and the iron-responsive GATA factor sequences, which came from *A. nidulans, Ustilago maydis, Phanerochate chrysosporium, Neurospora crassa, Candida albicans,* and *Schizosaccharomyces pombe*. Identification of *entA-entF* genes in bacterial genomes was performed using *E. coli* protein sequences as queries for BLASTP and TBLASTN to search all 1,382 Enterobacteriaceae genomes and proteomes downloaded from GenBank. Only genes from the 207 genomes where all six genes could be identified at E-value cutoff of 1E-10 were considered for further phylogenetic analyses.

#### Genome sequencing, assembly, and annotation

Yeast strains were obtained from the USDA Agricultural Research Service (ARS) NRRL Culture Collection in Peoria, Illinois, USA. Genomic DNA (gDNA) was isolated from individual strains, sonicated and ligated to Illumina sequencing adaptors as previously described (*56*). The paired-end libraries were submitted for 2×250bp sequencing on an Illumina HiSeq 2500 instrument. To generate whole-genome assemblies, Illumina reads were used as input to the meta-assembler pipeline iWGS v1.01 (*57*). Briefly, this pipeline performed quality-based read trimming, followed by k-mer length optimization, and used a range of state-of-the-art assemblers to generate multiple genome assemblies. Assembly quality was assessed using QUAST v4.4 (*58*), and the best assembly for each species was chosen based on the N_50_ statistic. Open reading frames were annotated in genomes using the MAKER pipeline v2 (*59*) and the GeneMark-ES v4.10 (*60*), Augustus v3.2.1 (*61*), and SNAP (release 2006-07-28) (*62*) gene predictors.

#### Phylogenetic reconstruction and topology tests

The species phylogeny was obtained by analyzing conserved single-copy fungal orthologs by using a previously described phylogenomic approach (*63*). Briefly, sequences of conserved, single-copy orthologous genes were identified in the genome assemblies using the BUSCO v3 software (*64*), single-copy BUSCO genes shared by at least 80% of species were aligned using MAFFT v7 (*65*), and these orthologs were used for maximum-likelihood phylogenetic reconstruction with RAxML v8 (*66*). The reconstruction was performed under the LG model of amino acid substitution (*67*) with empirical amino acid frequencies, four gamma distribution rate categories to estimate rate heterogeneity, and 100 rapid bootstrap pseudoreplicates. A concatenated super-alignment of all genes was also used for phylogenetic reconstruction by running ExaML v3.0.18 (*68*) under the JTT substitution matrix (chosen by the built-in maximum-likelihood model selection), per-site rate heterogeneity model with median approximation of the GAMMA rates, and with memory saving option for gappy alignments turned on. Constrained phylogeny reconstructions were conducted in RAxML through the “-g” option, and the AU topology tests were performed with IQ-TREE v1.5.4 (*69*) using 10,000 bootstrap pseudoreplicates.

Three evolutionary scenarios were considered to explain the course of the horizontal transfer event: (I) single-source, single-target; (II) single-source, multiple-targets; (III) and multiple-sources. Scenario I predicted that the yeast sequences would form a strongly-supported monophyletic group with a consistent internal topology. Scenario II predicted that yeast sequences would form a strongly-supported monophyletic group but not follow a consistent internal topology. Scenario III predicted that yeast sequences would not form a monophyletic group.

#### RNA sequencing and transcriptomics analyses

Cells were grown in quadruplicates for either 3 or 6 days on YPD agar, and RNA was extracted using the hot acid phenol protocol (*70*). Extracts were then treated with DNase to remove any residual DNA prior to treatment with the RNA Clean & Concentrator kit (Zymo Research #R1017, R1018). Total RNA yields were quantified with the Qubit RNA Assay Kit (Thermo Fisher). Next, mRNA was isolated and converted to cDNA using the NEBNext Poly(A) mRNA Magnetic Isolation Module (NEB #E7490) and prepared into Illumina libraries using the NEBNext Multiplex Oligos for Illumina (New England Biolabs #E7335, E7500). Library quality was assessed by gel electrophoresis and with the Qubit dsDNA Kit (Thermo Fisher) prior to submission for 2×125 paired-end sequencing with an Illumina HiSeq 2500 instrument. Reads were mapped to their respective genome assemblies using GSNAP (*71*) from the GMAP package (release date 2017-05-08) with the novel splicing site search option enabled. *De novo* transcriptome assembly was performed using the Trinity pipeline v2.4.0 (*72*), which was run in the RF strand-specific mode with the jaccard-clip option enabled. Transcript abundance of siderophore biosynthesis genes were estimated using StringTie v1.3.3b (*73*).

Evidence of transcriptional processing was evaluated by inspecting parts of the RNA-Seq reads that were soft-clipped from the ends of reads during the mapping step. 3’ ends were inspected for evidence of poly(A) tails of at least three consecutive As or Ts, which were not encoded in the genome. The power of such analysis is limited by the fact that only small fraction of reads (∼0.05%) are expected to be initiated using the (A)_6_ or (T)_6_ primers, which increases the rate of false negative results, but true positive results remain unaffected. With the above caveat, we note that evidence of poly(A) tails was not detected from the *C. versatilis entE, entA,* and *entH* genes. 5’ ends were inspected for presence of common sequences, encoded elsewhere in the genome, which could have been indicative of splicing leaders (in case of trans-splicing) or casette exons (in case of alternative cis-splicing).

#### Microbial culturing and chromeazurol S overlay (O-CAS) assays

Low-iron synthetic complete (SC) medium consisted of 5 g/L ammonium sulfate, 1.7 g/L Yeast Nitrogen Base (without amino acids, carbohydrates, ammonium sulfate, ferric chloride, or cupric sulfate), 2 g/L complete dropout mix, 2% dextrose (added after autoclaving), and 200 nM cupric sulfate. M9 minimal medium consisted of 0.4% glucose, 2 mM magnesium sulfate, 100 µM calcium dichloride, and 1x M9 salts (added as a 5x stock solution consisting of 64g/L dibasic sodium phosphate heptahydrate, 15 g/L monobasic potassium phosphate, 2.5 g/L sodium chloride, and 5 g/L ammonium chloride in deionized water).

The O-CAS Assay was carried out as previously described (*74*), with some modifications. Specifically, 10X CAS Blue Dye was made by combining the following: 50 mL Solution 1: (60 mg chromeazurol S dissolved in 50 mL deionized H_2_O), 9 mL Solution 2: (13.5 mg ferric chloride hexahydrate dissolved in 50 mL 10 mM hydrochloric acid) and 40 mL Solution 3: (73 mg hexadecyltrimethylammonium bromide (HDTMA) in 40 mL deionized H_2_O). Separately, 15.12 g PIPES (free acid) was added to 425 mL deionized water and adjusted to a pH of approximately 6.8 with 2.46 g sodium hydroxide. 4.5 g agarose was added as a solidifying agent, and the resulting solution was brought up to 450 mL with deionized water in a 1-L Erlenmeyer flask. To make the CAS overlay, the agarose-PIPES solution was heated to melt the agarose and added in a ratio of 9:1 to 10X CAS Blue Dye, and 6 mL of the resulting O-CAS solution were overlaid onto low-iron SC plates.

Yeast strains were grown to saturation in 3 mL YPD medium at 30 degrees centigrade on a rotating culture wheel, centrifuged at 3000 rpm for 5 minutes to collect the cells, and resuspended in 3 mL deionized water. A volume of 5 µL of the resulting cell suspension was spotted onto 60 mm diameter plates containing low-iron SC medium using agarose (1% w/v) as a gelling agent and incubated at 30 degrees centigrade for 7 days before adding 6 mL of O-CAS solution. *E. coli* cells were grown overnight in M9 minimal medium at 37 degrees centigrade, and 5 µL of culture was spotted onto low-iron SC plates that had already been overlaid with 6 mL of O-CAS solution and allowed to dry for at least 1 hour. Pictures of yeast colonies were taken 2 days after the O-CAS was poured, while *E. coli* colonies were photographed 5 days after the O-CAS was poured. With exposure and focus lock enabled, pictures were taken of the plates set on top of a miniature white light trans-illuminator placed under a gel-imaging dark box.

**Fig. S1.**
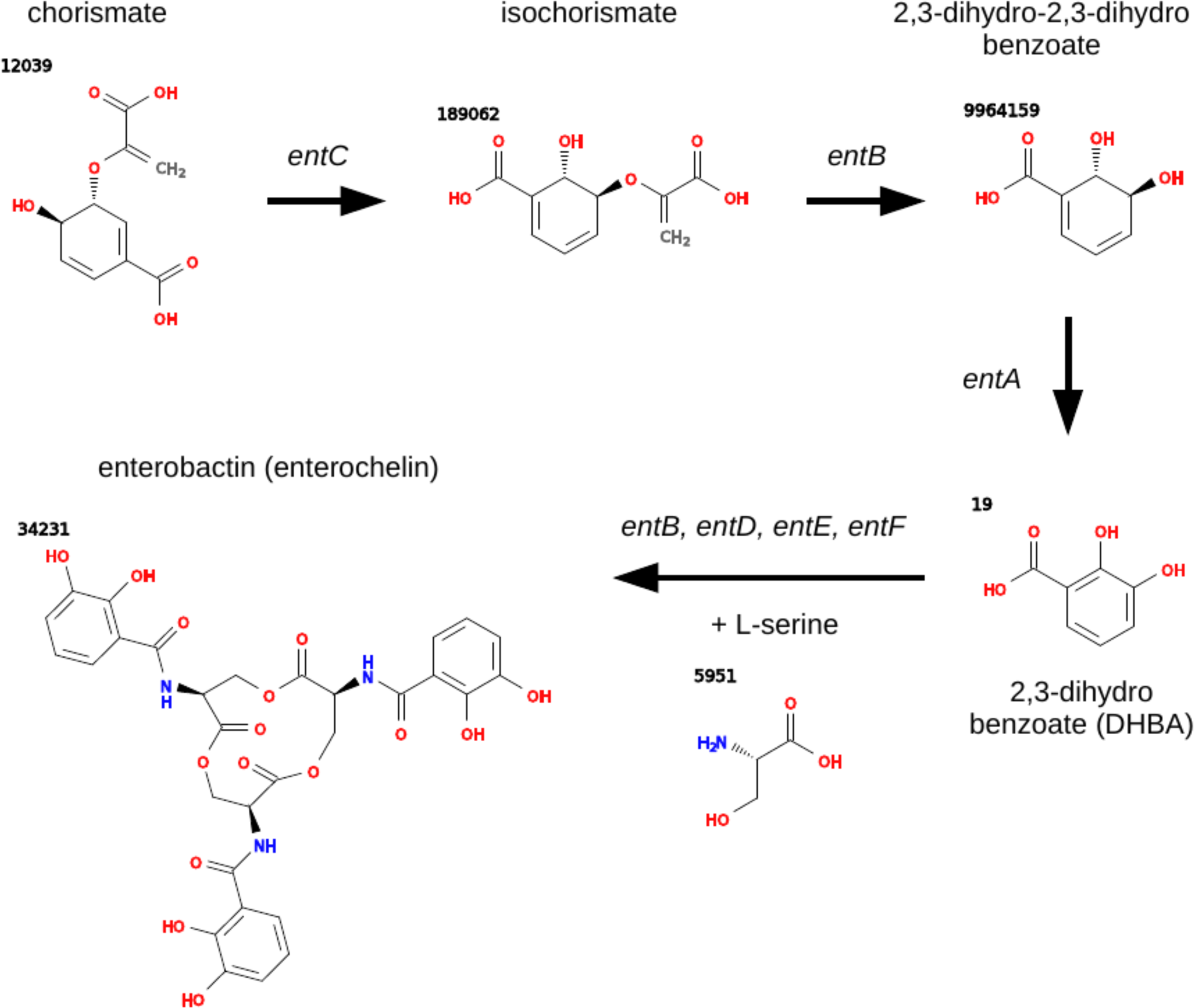
Enterobactin biosynthesis from chorismate. Genes involved at each biosynthesis step are marked above arrows. Intermediates of the final biosynthesis step have not yet been determined. PubChem IDs of individual chemical structures are marked in bold. Chorismate biosynthesis is broadly conserved because it is a key compound in the shikimate pathway that is involved in biosynthesis of aromatic amino acids. *entH* encodes a proofreading thioesterase whose activity is not required for the production of enterobactin and related catecholate-class siderophores.

**Fig. S2.**
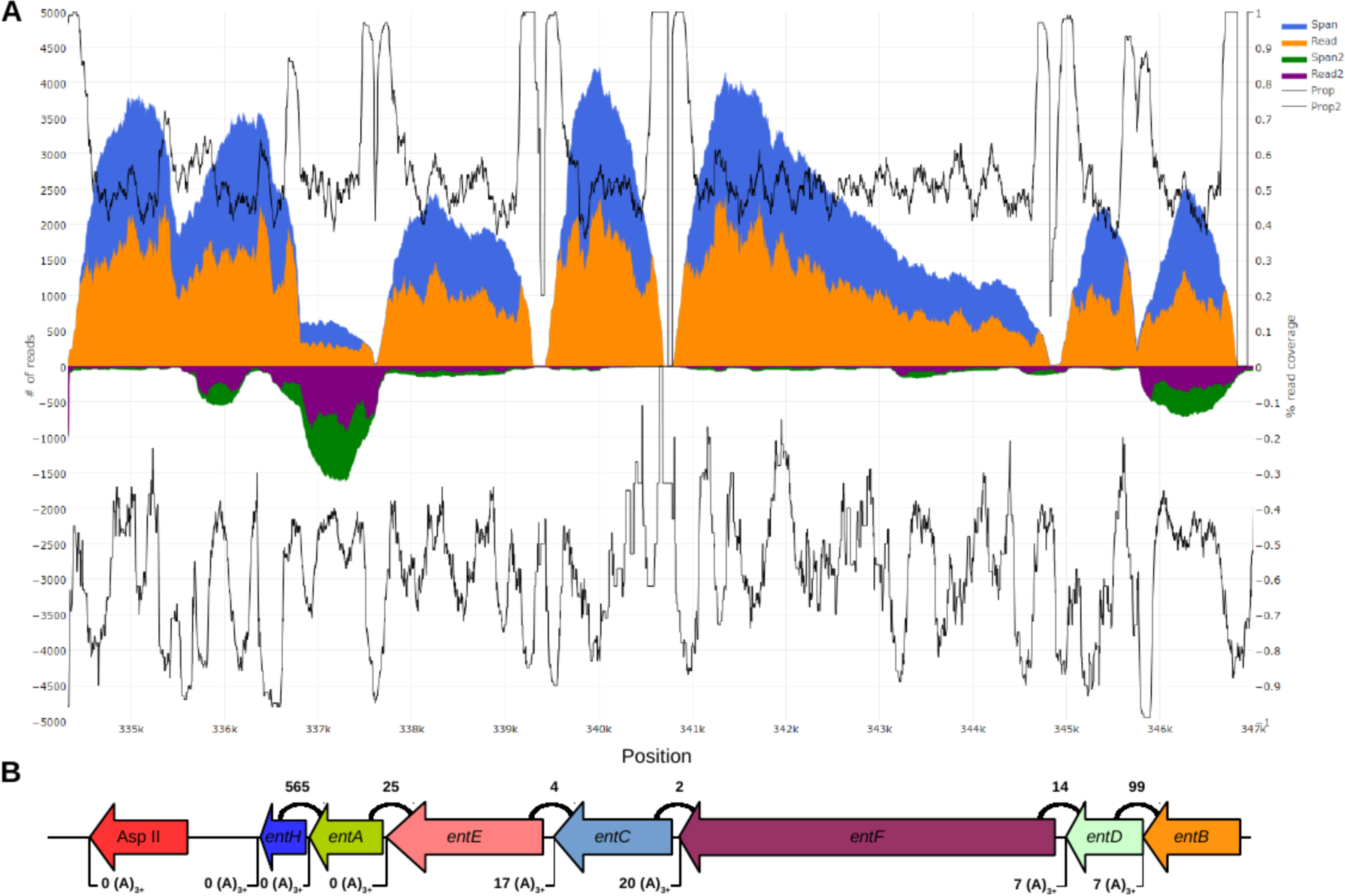
Transcriptomics of the siderophore biosynthesis genes in *C. versatilis*. (A) The orange area indicates per-base coverage by RNA-seq reads (read coverage). The blue area indicates per-base cumulative coverage by RNA-seq reads and inserts between read pairs (span coverage). The purple area indicates read coverage of the opposite strand. The green area indicates span coverage of the opposite strand. The black lines indicate the ratios of the read coverage over the span coverage data for the relevant strand. (B) Diagram of siderophore biosynthesis genes in the *C. versatilis* genome, drawn to scale, including a bacterial class II asparaginase downstream from the operon. Counts above indicate read pairs cross-mapping between genes. Counts below indicate reads containing putative poly(A) tails.

**Fig. S3.**
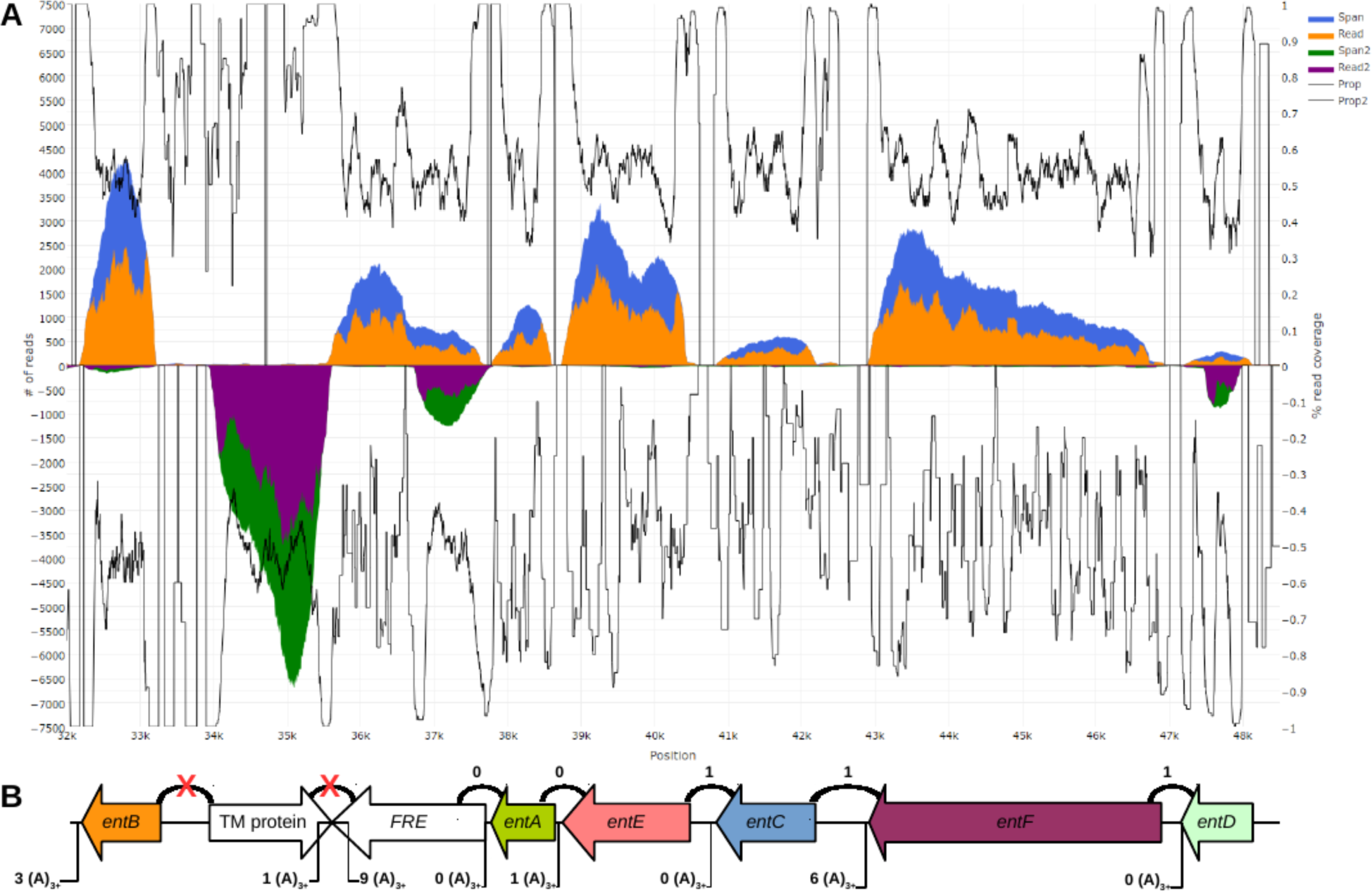
Transcriptomics of the siderophore biosynthesis genes in *St. bombicola.* (A) The orange area indicates per-base coverage by RNA-seq reads (read coverage). The blue area indicates per-base cumulative coverage by RNA-seq reads and inserts between read pairs (span coverage). The purple area indicates read coverage of the opposite strand. The green area indicates span coverage of the opposite strand. The black lines indicate the ratios of the read coverage over the span coverage data for the relevant strand. (B) Diagram of siderophore biosynthesis genes in the *St. bombicola* genome, drawn to scale. Counts above indicate read pairs cross-mapping between genes. Counts below indicate reads containing putative poly(A) tails.

**Fig. S4.**
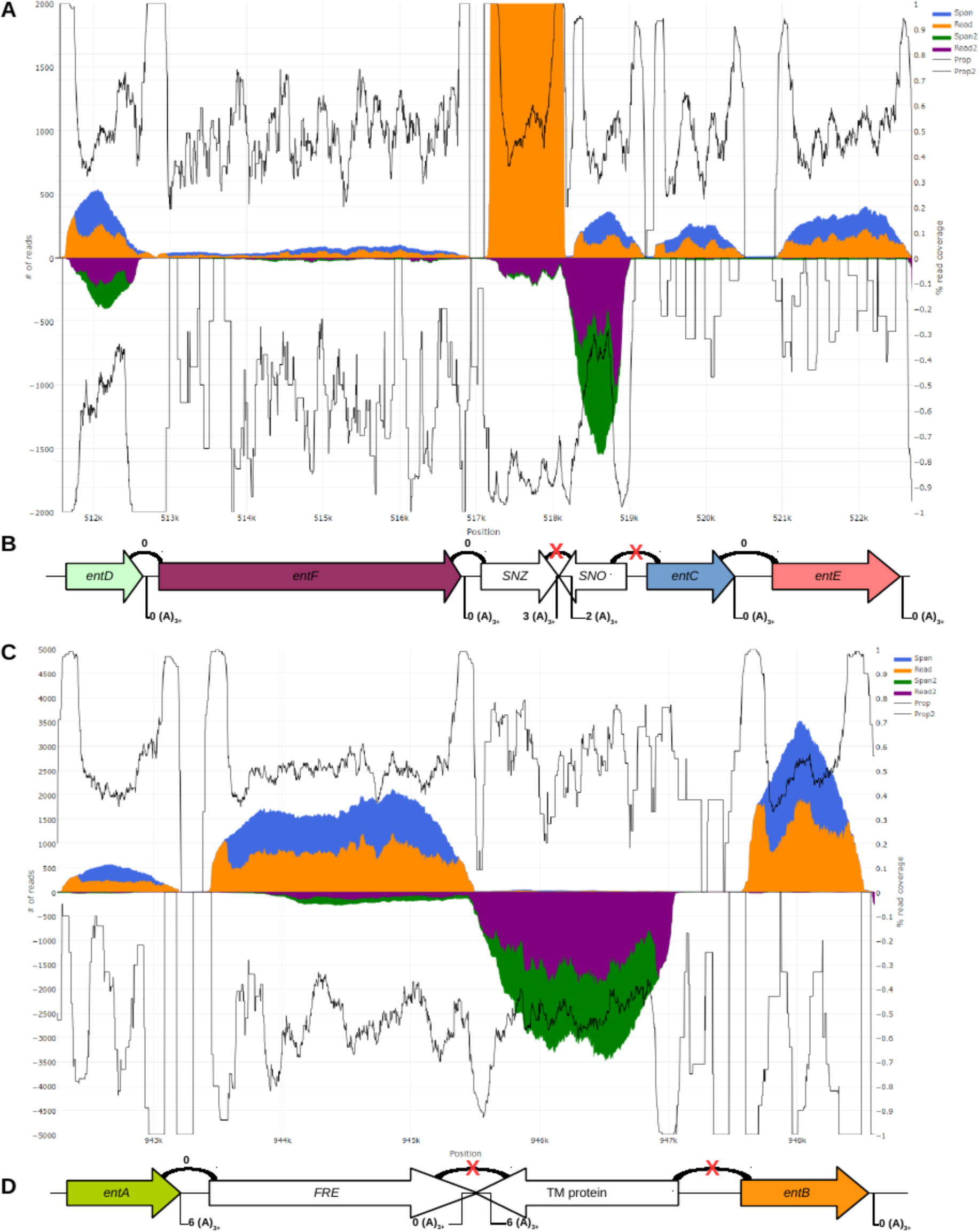
Transcriptomics of the siderophore biosynthesis genes in *C. apicola.* (A, C) The orange area indicates per-base coverage by RNA-seq reads (read coverage). The blue area indicates per-base cumulative coverage by RNA-seq reads and inserts between read pairs (span coverage). The purple area indicates read coverage of the opposite strand. The green area indicates span coverage of the opposite strand. The black lines indicate the ratios of the read coverage over the span coverage data for the relevant strand. (B, D) Diagrams of siderophore biosynthesis genes in the *C. apicola* genome, drawn to scale. Counts above indicate read pairs cross-mapping between genes. Counts below indicate reads containing putative poly(A) tails.

**Fig. S5.**
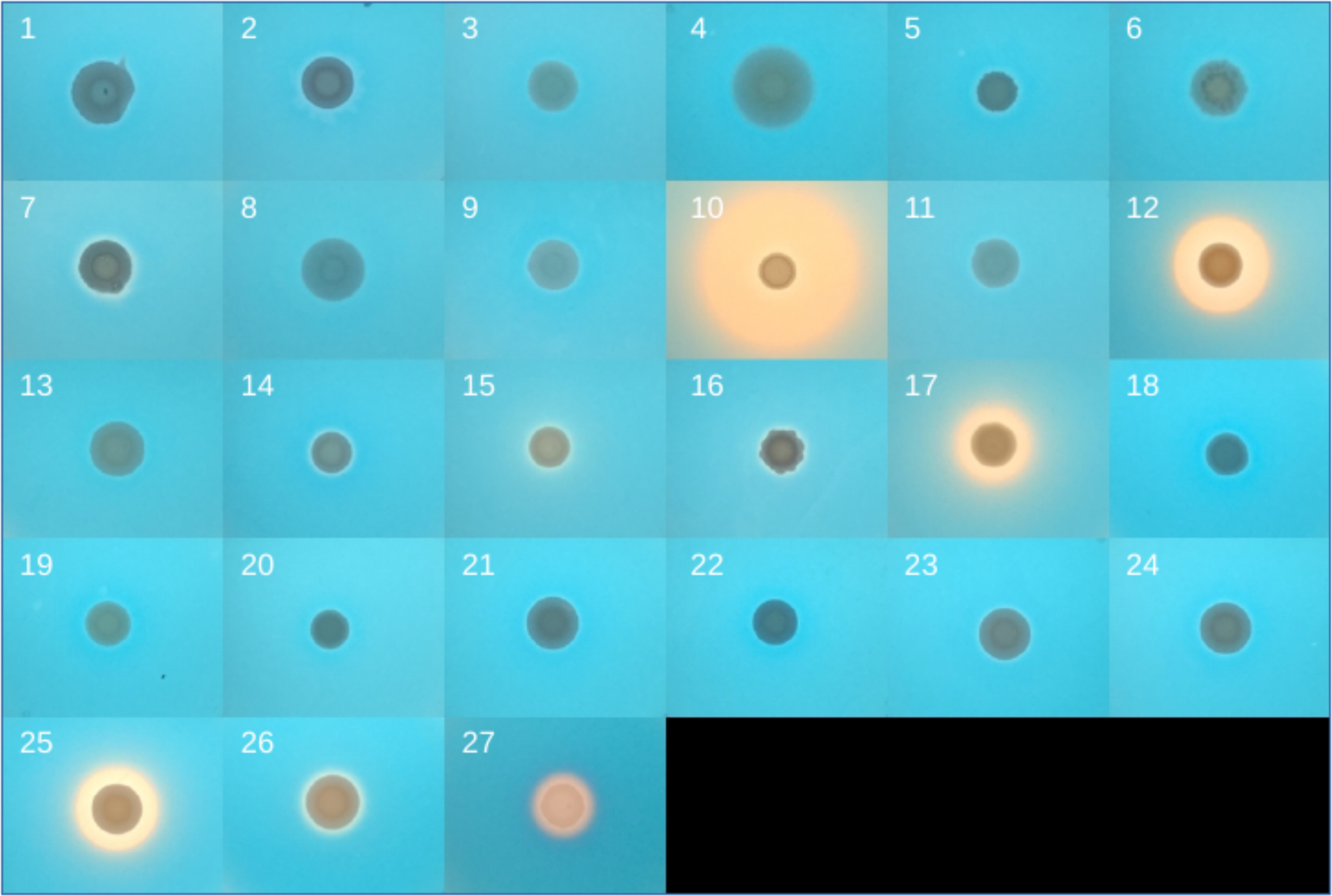
CAS assay for all species studied. Species legend: (1) *Saccharomyces cerevisiae* FM1282, (2) *Kluyveromyces lactis* CBS 2359, (3) *Tortispora caseinolytica* NRRL Y-17796^T^, (4) *Yarrowia keelungensis* NRRL Y-63742^T^, (5) *Yarrowia deformans* NRRL Y-321^T^, (6) *Yarrowia lipolytica* NRRL YB-423^T^, (7) *Blastobotrys (Arxula) adeninivorans* NRRL Y-17592, (8) *Sugiyamaella lignohabitans* NRRL YB-1473^T^, (9) *Candida hasegawae* JCM 12559^T^, (10) *Candida pararugosa* NRRL Y-17089^T^, (11) *Wickerhamiella cacticola* NRRL Y-27362^T^, (12) *Candida versatilis* NRRL Y-6652^T^, (13) *Candida gropengiesseri* NRRL Y-17142^T^, (14) *Candida davenportii* CBS 9069^T^, (15) *Candida ratchasimensis* CBS 10611^T^, (16) *Candida apicola* NRRL Y-2481^T^, (17) *Candida riodocensis* NRRL Y-27859^T^, (18) *Candida infanticola* NRRL Y-17858^T^, (19) *Wickerhamiella occidentalis* NRRL Y-27364, (20) *Wickerhamiella domercqiae* NRRL Y-6692^T^, (21) *Candida tilneyi* CBS 8794^T^, (22) *Candida sorbosivorans* CBS 8768^T^, (23) *Candida geochares* NRRL Y-17073^T^, (24) *Candida vaccinii* NRRL Y-17684^T^, (25) *Candida kuoi* NRRL Y-27208^T^, (26) *Starmerella bombicola* NRRL Y-17069^T^, and (27) *Escherichia coli* MG1655 (positive control).

**Table S1 (separate file)**

List of genomes and putative hits to iron utilization proteins analyzed in this study. Unless otherwise specified, query proteins were from *S. cerevisiae.*

**Table S2 (separate file)**

List and statistics of novel yeast genomes sequenced in this study.

**Table S3 (separate file)**

List of Gammoproteobacteria species with the six siderophore biosynthesis genes (*entA-entF*) analyzed in this study.

**Table S4 (separate file)**

Newick format strings of phylogenetic trees reconstructed from individual *ent* genes and from the concatenated superalignments.

**Table S5 (separate file)**

P-values from the AU test between phylogenies reconstructed under 12-species and 11-species monophyletic constraints.

**Table S6 (separate file)**

Normalized per-base coverage of the *ent* genes from *C. versatilis, C. apicola,* and *St. bombicola.*

## References and Notes

1. T. Blumenthal, K. S. Gleason, *Caenorhabditis elegans* operons: form and function. Nat. Rev. Genet. 4, 112–120 (2003).

2. J. Spieth, G. Brooke, S. Kuersten, K. Lea, T. Blumenthal, Operons in *C. elegans*: polycistronic mRNA precursors are processed by trans-splicing of SL2 to downstream coding regions. Cell. 73, 521–532 (1993).

3. A. E. Vandenberghe, T. H. Meedel, K. E. M. Hastings, mRNA 5′-leader trans-splicing in the chordates. Genes Dev. 15, 294–303 (2001).

4. P. Ganot, T. Kallesøe, R. Reinhardt, D. Chourrout, E. M. Thompson, Spliced-Leader RNA trans Splicing in a Chordate, *Oikopleura dioica*, with a Compact Genome. Mol. Cell. Biol. 24, 7795–7805 (2004).

5. M. V. Omelchenko, K. S. Makarova, Y. I. Wolf, I. B. Rogozin, E. V. Koonin, Evolution of mosaic operons by horizontal gene transfer and gene displacement in situ. Genome Biol. 4, R55 (2003).

6. J. G. Lawrence, J. R. Roth, Selfish Operons: Horizontal Transfer May Drive the Evolution of Gene Clusters. Genetics. 143, 1843–1860 (1996).

7. P. J. Keeling, J. D. Palmer, Horizontal gene transfer in eukaryotic evolution. Nat. Rev. Genet. 9, 605–618 (2008).

8. S. M. Soucy, J. Huang, J. P. Gogarten, Horizontal gene transfer: building the web of life. Nat. Rev. Genet. 16, 472–482 (2015).

9. W. G. Alexander, J. H. Wisecaver, A. Rokas, C. T. Hittinger, Horizontally acquired genes in early-diverging pathogenic fungi enable the use of host nucleosides and nucleotides. Proc. Natl. Acad. Sci. U. S. A. 113, 4116–4121 (2016).

10. T. A. Richards, D. M. Soanes, M. D. M. Jones, O. Vasieva, G. Leonard, K. Paszkiewicz, P. G. Foster, N. Hall, N. J. Talbot, Horizontal gene transfer facilitated the evolution of plant parasitic mechanisms in the oomycetes. Proc. Natl. Acad. Sci. U. S. A. 108, 15258–15263 (2011).

11. J. C. Slot, A. Rokas, Horizontal Transfer of a Large and Highly Toxic Secondary Metabolic Gene Cluster between Fungi. Curr. Biol. 21, 134–139 (2011).

12. S. C. Andrews, A. K. Robinson, F. Rodríguez-Quiñones, Bacterial iron homeostasis. FEMS Microbiol. Rev. 27, 215–237 (2003).

13. A. Sheftel, O. Stehling, R. Lill, Iron–sulfur proteins in health and disease. Trends Endocrinol. Metab. 21, 302–314 (2010).

14. R. Sutak, E. Lesuisse, J. Tachezy, D. R. Richardson, Crusade for iron: iron uptake in unicellular eukaryotes and its significance for virulence. Trends Microbiol. 16, 261–268 (2008).

15. D. H. Scharf, T. Heinekamp, A. A. Brakhage, Human and Plant Fungal Pathogens: The Role of Secondary Metabolites. PLOS Pathog. 10, e1003859 (2014).

16. E. P. Skaar, The battle for iron between bacterial pathogens and their vertebrate hosts. PLoS Pathog. 6, e1000949 (2010).

17. I. K. Toth, L. Pritchard, P. R. J. Birch, Comparative Genomics Reveals What Makes An Enterobacterial Plant Pathogen. Annu. Rev. Phytopathol. 44, 305–336 (2006).

18. C. Wandersman, P. Delepelaire, Bacterial Iron Sources: From Siderophores to Hemophores. Annu. Rev. Microbiol. 58, 611–647 (2004).

19. H. Haas, M. Eisendle, B. G. Turgeon, Siderophores in fungal physiology and virulence. Annu. Rev. Phytopathol. 46, 149–187 (2008).

20. M. Gilliam, Identification and roles of non-pathogenic microflora associated with honey bees. FEMS Microbiol. Lett. 155, 1–10 (1997).

21. C. T. Walsh, J. Liu, F. Rusnak, M. Sakaitani, Molecular studies on enzymes in chorismate metabolism and the enterobactin biosynthetic pathway. Chem. Rev. 90, 1105–1129 (1990).

22. A. Levin-Karp, U. Barenholz, T. Bareia, M. Dayagi, L. Zelcbuch, N. Antonovsky, E. Noor, R. Milo, Quantifying Translational Coupling in E. coli Synthetic Operons Using RBS Modulation and Fluorescent Reporters. ACS Synth. Biol. 2, 327–336 (2013).

23. S. Okuda, S. Kawashima, K. Kobayashi, N. Ogasawara, M. Kanehisa, S. Goto, Characterization of relationships between transcriptional units and operon structures in *Bacillus subtilis* and *Escherichia coli*. BMC Genomics. 8, 48 (2007).

24. L. David, W. Huber, M. Granovskaia, J. Toedling, C. J. Palm, L. Bofkin, T. Jones, R. W. Davis, L. M. Steinmetz, A high-resolution map of transcription in the yeast genome. Proc. Natl. Acad. Sci. 103, 5320–5325 (2006).

25. V. Pelechano, W. Wei, L. M. Steinmetz, Extensive transcriptional heterogeneity revealed by isoform profiling. Nature. 497, 127–131 (2013).

26. K. V. Doren, D. Hirsh, mRNAs that mature through trans-splicing in *Caenorhabditis elegans* have a trimethylguanosine cap at their 5’ termini. Mol. Cell. Biol. 10, 1769–1772 (1990).

27. H. Keren, G. Lev-Maor, G. Ast, Alternative splicing and evolution: diversification, exon definition and function. Nat. Rev. Genet. 11, 345–355 (2010).

28. N. Khaldi, J. Collemare, M.-H. Lebrun, K. H. Wolfe, Evidence for horizontal transfer of a secondary metabolite gene cluster between fungi. Genome Biol. 9, R18 (2008).

29. J. C. Slot, A. Rokas, Multiple GAL pathway gene clusters evolved independently and by different mechanisms in fungi. Proc. Natl. Acad. Sci. U. S. A. 107, 10136–10141 (2010).

30. M. A. Campbell, M. Staats, J. A. L. van Kan, A. Rokas, J. C. Slot, Repeated loss of an anciently horizontally transferred gene cluster in *Botrytis*. Mycologia. 105, 1126–1134 (2013).

31. R. H. Proctor, F. Van Hove, A. Susca, G. Stea, M. Busman, T. van der Lee, C. Waalwijk, A. Moretti, T. J. Ward, Birth, death and horizontal transfer of the fumonisin biosynthetic gene cluster during the evolutionary diversification of *Fusarium*. Mol. Microbiol. 90, 290–306 (2013).

32. D. Leduc, A. Battesti, E. Bouveret, The hotdog thioesterase EntH (YbdB) plays a role in vivo in optimal enterobactin biosynthesis by interacting with the ArCP domain of EntB. J. Bacteriol. 189, 7112–7126 (2007).

33. J. W. Taylor, M. L. Berbee, Dating divergences in the Fungal Tree of Life: review and new analyses. Mycologia. 98, 838–849 (2006).

34. D. A. Fitzpatrick, Horizontal gene transfer in fungi. FEMS Microbiol. Lett. 329, 1–8 (2012).

35. J. C. D. Hotopp, M. E. Clark, D. C. S. G. Oliveira, J. M. Foster, P. Fischer, M. C. M. Torres, J. D. Giebel, N. Kumar, N. Ishmael, S. Wang, J. Ingram, R. V. Nene, J. Shepard, J. Tomkins, S. Richards, D. J. Spiro, E. Ghedin, B. E. Slatko, H. Tettelin, J. H. Werren, Widespread Lateral Gene Transfer from Intracellular Bacteria to Multicellular Eukaryotes. Science. 317, 1753–1756 (2007).

36. M. Marcet-Houben, T. Gabaldón, Acquisition of prokaryotic genes by fungal genomes. Trends Genet. 26, 5–8 (2010).

37. T. A. Richards, J. B. Dacks, J. M. Jenkinson, C. R. Thornton, N. J. Talbot, Evolution of Filamentous Plant Pathogens: Gene Exchange across Eukaryotic Kingdoms. Curr. Biol. 16, 1857–1864 (2006).

38. B. J. Weigel, S. G. Burgett, V. J. Chen, P. L. Skatrud, C. A. Frolik, S. W. Queener, T. D. Ingolia, Cloning and expression in *Escherichia coli* of isopenicillin N synthetase genes from *Streptomyces lipmanii* and *Aspergillus nidulans*. J. Bacteriol. 170, 3817–3826 (1988).

39. J. H. Wisecaver, A. Rokas, Fungal metabolic gene clusters-caravans traveling across genomes and environments. Front. Microbiol. 6, 161 (2015).

40. T. A. Richards, G. Leonard, D. M. Soanes, N. J. Talbot, Gene transfer into the fungi. Fungal Biol. Rev. 25, 98–110 (2011).

41. A. Routh, T. Domitrovic, J. E. Johnson, Host RNAs, including transposons, are encapsidated by a eukaryotic single-stranded RNA virus. Proc. Natl. Acad. Sci. 109, 1907–1912 (2012).

42. J.-F. Flot, B. Hespeels, X. Li, B. Noel, I. Arkhipova, E. G. J. Danchin, A. Hejnol, B. Henrissat, R. Koszul, J.-M. Aury, V. Barbe, R.-M. Barthélémy, J. Bast, G. A. Bazykin, O. Chabrol, A. Couloux, M. Da Rocha, C. Da Silva, E. Gladyshev, P. Gouret, O. Hallatschek, B. Hecox-Lea, K. Labadie, B. Lejeune, O. Piskurek, J. Poulain, F. Rodriguez, J. F. Ryan, O. A. Vakhrusheva, E. Wajnberg, B. Wirth, I. Yushenova, M. Kellis, A. S. Kondrashov, D. B. Mark Welch, P. Pontarotti, J. Weissenbach, P. Wincker, O. Jaillon, K. Van Doninck, Genomic evidence for ameiotic evolution in the bdelloid rotifer *Adineta vaga*. Nature. 500, 453–457 (2013).

43. E. A. Gladyshev, M. Meselson, I. R. Arkhipova, Massive Horizontal Gene Transfer in Bdelloid Rotifers. Science. 320, 1210–1213 (2008).

44. W. F. Doolittle, You are what you eat: a gene transfer ratchet could account for bacterial genes in eukaryotic nuclear genomes. Trends Genet. 14, 307–311 (1998).

45. M.-A. Lachance, W. T. Starmer, C. A. Rosa, J. M. Bowles, J. S. F. Barker, D. H. Janzen, Biogeography of the yeasts of ephemeral flowers and their insects. FEMS Yeast Res. 1, 1–8 (2001).

46. C. A. Rosa, M.-A. Lachance, The yeast genus *Starmerella* gen. nov. and *Starmerella bombicola* sp. nov., the teleomorph of *Candida bombicola* (Spencer, Gorin & Tullock) Meyer & Yarrow. Int. J. Syst. Evol. Microbiol. 48, 1413–1417 (1998).

47. C. A. Rosa, M. A. Lachance, J. O. C. Silva, A. C. P. Teixeira, M. M. Marini, Y. Antonini, R. P. Martins, Yeast communities associated with stingless bees. FEMS Yeast Res. 4, 271–275 (2003).

48. K. Watanabe, M. Sato, Plasmid-Mediated Gene Transfer Between Insect-Resident Bacteria, *Enterobacter cloacae*, and Plant-Epiphytic Bacteria, *Erwinia herbicola*, in Guts of Silkworm Larvae. Curr. Microbiol. 37, 352–355 (1998).

49. K. Watanabe, W. Hara, M. Sato, Evidence for Growth of Strains of the Plant Epiphytic Bacterium *Erwinia herbicola* and Transconjugation among the Bacterial Strains in Guts of the Silkworm *Bombyx mori*. J Invertebr Pathol. 72, 104–111 (1998).

50. I. Stefanini, L. Dapporto, L. Berná, M. Polsinelli, S. Turillazzi, D. Cavalieri, Social wasps are a *Saccharomyces* mating nest. Proc. Natl. Acad. Sci. 113, 2247–2251 (2016).

51. J. A. Heinemann, G. F. Sprague, Bacterial conjugative plasmids mobilize DNA transfer between bacteria and yeast. Nature. 340, 205–209 (1989).

52. M. F. Barber, N. C. Elde, Buried Treasure: Evolutionary Perspectives on Microbial Iron Piracy. Trends Genet. 31, 627–636 (2015).

53. Z. Chen, K. A. Lewis, R. K. Shultzaberger, I. G. Lyakhov, M. Zheng, B. Doan, G. Storz, T. D. Schneider, Discovery of Fur binding site clusters in *Escherichia coli* by information theory models. Nucleic Acids Res. 35, 6762–6777 (2007).

54. H. Boukhalfa, A. L. Crumbliss, Chemical aspects of siderophore mediated iron transport. Biometals. 15, 325–339 (2002).

55. S. F. Altschul, W. Gish, W. Miller, E. W. Myers, D. J. Lipman, Basic local alignment search tool. J. Mol. Biol. 215, 403–410 (1990).

56. C. T. Hittinger, P. Gonçalves, J. P. Sampaio, J. Dover, M. Johnston, A. Rokas, Remarkably ancient balanced polymorphisms in a multi-locus gene network. Nature. 464, 54–58 (2010).

57. X. Zhou, D. Peris, J. Kominek, C. P. Kurtzman, C. T. Hittinger, A. Rokas, in silico Whole Genome Sequencer & Analyzer (iWGS): A Computational Pipeline to Guide the Design and Analysis of de novo Genome Sequencing Studies. G3. 6, 3655–3662 (2016).

58. A. Gurevich, V. Saveliev, N. Vyahhi, G. Tesler, QUAST: quality assessment tool for genome assemblies. Bioinformatics. 29, 1072–1075 (2013).

59. C. Holt, M. Yandell, MAKER2: an annotation pipeline and genome-database management tool for second-generation genome projects. BMC Bioinformatics. 12, 491 (2011).

60. V. Ter-Hovhannisyan, A. Lomsadze, Y. O. Chernoff, M. Borodovsky, Gene prediction in novel fungal genomes using an ab initio algorithm with unsupervised training. Genome Res. 18, 1979–1990 (2008).

61. M. Stanke, M. Diekhans, R. Baertsch, D. Haussler, Using native and syntenically mapped cDNA alignments to improve de novo gene finding. Bioinformatics. 24, 637–644 (2008).

62. I. Korf, Gene finding in novel genomes. BMC Bioinformatics. 5, 59 (2004).

63. X.-X. Shen, X. Zhou, J. Kominek, C. P. Kurtzman, C. T. Hittinger, A. Rokas, Reconstructing the Backbone of the Saccharomycotina Yeast Phylogeny Using Genome-Scale Data. G3. 6, 3927–3939 (2016).

64. F. A. Simão, R. M. Waterhouse, P. Ioannidis, E. V. Kriventseva, E. M. Zdobnov, BUSCO: assessing genome assembly and annotation completeness with single-copy orthologs. Bioinformatics. 31, 3210–3212 (2015).

65. K. Katoh, D. M. Standley, MAFFT multiple sequence alignment software version 7: improvements in performance and usability. Mol. Biol. Evol. 30, 772–780 (2013).

66. A. Stamatakis, RAxML version 8: a tool for phylogenetic analysis and post-analysis of large phylogenies. Bioinformatics. 30, 1312–1313 (2014).

67. S. Q. Le, O. Gascuel, An improved general amino acid replacement matrix. Mol. Biol. Evol. 25, 1307–1320 (2008).

68. A. M. Kozlov, A. J. Aberer, A. Stamatakis, ExaML version 3: a tool for phylogenomic analyses on supercomputers. Bioinformatics. 31, 2577–2579 (2015).

69. L.-T. Nguyen, H. A. Schmidt, A. von Haeseler, B. Q. Minh, IQ-TREE: a fast and effective stochastic algorithm for estimating maximum-likelihood phylogenies. Mol. Biol. Evol. 32, 268–274 (2015).

70. P. Chomczynski, N. Sacchi, Single-step method of RNA isolation by acid guanidinium thiocyanate-phenol-chloroform extraction. Anal. Biochem. 162, 156–159 (1987).

71. T. D. Wu, S. Nacu, Fast and SNP-tolerant detection of complex variants and splicing in short reads. Bioinformatics. 26, 873–881 (2010).

72. M. G. Grabherr, B. J. Haas, M. Yassour, J. Z. Levin, D. A. Thompson, I. Amit, X. Adiconis, L. Fan, R. Raychowdhury, Q. Zeng, Z. Chen, E. Mauceli, N. Hacohen, A. Gnirke, N. Rhind, F. di Palma, B. W. Birren, C. Nusbaum, K. Lindblad-Toh, N. Friedman, A. Regev, Full-length transcriptome assembly from RNA-Seq data without a reference genome. Nat. Biotechnol. 29, 644–652 (2011).

73. M. Pertea, G. M. Pertea, C. M. Antonescu, T.-C. Chang, J. T. Mendell, S. L. Salzberg, StringTie enables improved reconstruction of a transcriptome from RNA-seq reads. Nat. Biotechnol. 33, 290–295 (2015).

74. S. Pérez-Miranda, N. Cabirol, R. George-Téllez, L. S. Zamudio-Rivera, F. J. Fernández, O-CAS, a fast and universal method for siderophore detection. J. Microbiol. Methods. 70, 127–131 (2007).

